# A non-coding RNA network involved in KSHV tumorigenesis

**DOI:** 10.1101/2021.03.05.434134

**Authors:** Julián Naipauer, Martín E. García Solá, Daria Salyakina, Santas Rosario, Sion Williams, Omar Coso, Martín C. Abba, Enrique A. Mesri, Ezequiel Lacunza

## Abstract

Regulatory pathways involving non-coding RNAs (ncRNAs) such as microRNAs (miRNAs) and long non-coding RNAs (lncRNA) have gained great relevance due to their role in the control of gene expression modulation. Using RNA sequencing of KSHV Bac36 transfected mouse endothelial cells (mECK36) and tumors, we have analyzed the host and viral transcriptome to uncover the role lncRNA-miRNA-mRNA driven networks in KSHV tumorigenesis. The integration of the differentially expressed ncRNAs, with an exhaustive computational analysis of their experimentally supported targets, led us to dissect complex networks integrated by the cancer-related lncRNAs Malat, Neat1, H19, Meg3 and their associated miRNA-target pairs. These networks would modulate pathways related to KSHV pathogenesis, such as viral carcinogenesis, p53 signaling, RNA surveillance, and Cell cycle control. Finally, the ncRNA-mRNA analysis allowed us to develop signatures that can be used to an appropriate identification of druggable gene or networks defining relevant AIDS-KS therapeutic targets.

## INTRODUCTION

Non-coding RNAs (ncRNAs) are RNA transcripts that do not encode proteins and based on the length can be divided into two classes: small ncRNAs (sncRNAs), with transcripts shorter than 200 nucleotides, and long ncRNAs (lncRNAs), with transcripts longer than 200 nucleotides [Ponting et al., 2009] Regulatory pathways involving ncRNAs, such as microRNAs (miRNAs), belonging to the class of sncRNA, and lncRNAs have gained great relevance due to their role in the control of gene (mRNA) expression. Different modes of interactions between lncRNAs and miRNAs have been reported: miRNA decay of lncRNAs, lncRNAs competing with mRNAs to bind to miRNAs, lncRNAs competing with miRNAs to bind to mRNA, and lncRNAs being shortened to miRNAs [Li et al., 2016, Hu et al., 2018]. All these interactions regulate the expression levels of mRNAs and in turn affect core protein signals, resulting in changes in the physiological functions of cells.

Kaposi’s sarcoma (KS) is an AIDS-associated malignancy caused by the KS herpesvirus (KSHV). Despite the reduction of its incidence since the implementation of anti-retroviral therapy (ART), KS continues to be a global, difficult-to-treat health problem, in particular for ART-resistant forms [Cesarman et al., 2019, Dittmer et al., 2007]. KS is characterized by the proliferation of KSHV-infected spindle cells and profuse angiogenesis [Mesri et al., 2010].

The life cycle of KSHV has two well-defined phases: latent and lytic. In the latent phase, the virus expresses a few genes involved in viral persistence and host immune evasion. During the lytic phase which is triggered by environmental and/or physiological stimuli the viral genome replicates and new virions are formed [Yan et al., 2019]. At this stage, KSHV is particularly effective at exploiting host gene expression for its own benefit. In this sense, the coevolution of the virus and its host has developed an intricate association between the virus genome, with its coding genes and non-coding genes, and the host RNA biosynthesis machinery [Macveigh et al., 2020]. To the point that KSHV can seize control of RNA surveillance pathways, such as DNA damage response (DDR), pre-mRNA control machinery and the Nonsense-mediated mRNA decay (NMD), to fine-tuningthe global gene expression environment throughout both phases of infection [Yan et al., 2019, Ohsaki et al., 2020, Zhao et al., 2020].

A recent study of KSHV-infected TIVE cells using wild-type and miRNA-deleted KSHV in conjunction with microarray technology to profile lncRNA expression found that KSHV can deregulate hundreds of host lncRNAs. These data established that KSHV de-regulates lncRNA in a miRNA-dependent fashion [Sethuraman et al., 2017].

Using deep RNA sequencing of KSHV Bac36 transfected mouse endothelial cells (mECK36) and tumors [Mutlu et al., 2007], we have previously analyzed the host and viral transcriptome to characterize mechanisms of KSHV dependent and independent sarcomagenesis as well as the contribution of host mutations [Naipauer et al., 2020]. We now decided to study in this model, in a genome-wide fashion, the ncRNAs landscape to better understand the relationship between mRNAs, lncRNAs and miRNAs in shaping KSHV tumorigenesis mechanisms.

This study allowed us to identify the most relevant lncRNAs involved in KSHV tumorigenesis through the mouse KS-model (*Malat1, Neat1, H19 and MEG3*). In addition to having common target genes, pathway analysis showed that the four lncRNAs also share common related processes, mainly associated with cancer and viral infections, which would contribute with a network of gene-pathways closely associated with KSHV oncogenesis.

On the other hand, small RNA-sequencing and miRNA analysis revealed a high proportion of upregulated host miRNAs dependent of KSHV infection, indicating that the presence of KSHV has a significant impact on the metabolism of host miRNAs, whose target genes are mainly associated to Angiogenesis, ECM, Spliceosome, p53 signaling, Viral infections and Cell cycle control. Similarly, functional analysis of KSHV miRNA targets showed enrichment in processes such as Cell cycle, Spliceosome, RNA transport, MicroRNA Regulation of DDR and p53 signaling, suggesting that viral miRNAs might mimic cellular miRNAs.

The integrative analysis of viral and host non-coding and coding RNAs and the related processes showed a landscape of the potential relationships of lncRNA-miRNA-mRNA in a KSHV setting. This network highlights that the upregulated genes are involved in processes previously related to KSHV tumorigenesis while downregulated genes are associated with host cell cycle checkpoints and RNA surveillance pathways: *G1 to S cycle control, p53 activity regulation, MicroRNA regulation of DDR, Spliceosome, RNA transport, E2F transcription factor network*. Finally, the ncRNA-mRNA analysis in the animal model presented here allowed us to develop signatures that can be used to identify druggable gene or networks defining relevant AIDS-KS therapeutic targets.

## RESULTS

### 1- Genome-wide analysis of Non-coding RNAs in a cell and animal model of Kaposi’s Sarcoma

To analyze the ncRNA expression profile in a cell and animal model of Kaposi’s Sarcoma we performed deep RNA-seq analysis of all the stages of this model. Figure 1A shows a schematic representation of the model: tumors formed by KSHV Bac36 transfected mouse endothelial cells, KSHV (+) cells, are all episomally infected with KSHV Bac36, and when KSHV (+) cells prior to form tumors lose the KSHV episome *in vitro* by withdrawal of antibiotic selection, KSHV (-) cells, they completely lose tumorigenicity [Mutlu et al., 2007, Naipauer et al., 2020]. In contrast to KSHV (-) cells, cells explanted from KSHV (+) BAC36 tumors and grown in the absence of antibiotic selection lose the KSHV episome, KSHV (-) tumor cells, are tumorigenic and are able to form KSHV (-) tumors [Mutlu et al., 2007; Naipauer et al., 2020; Ma et al., 2013].

**Figure 1.**
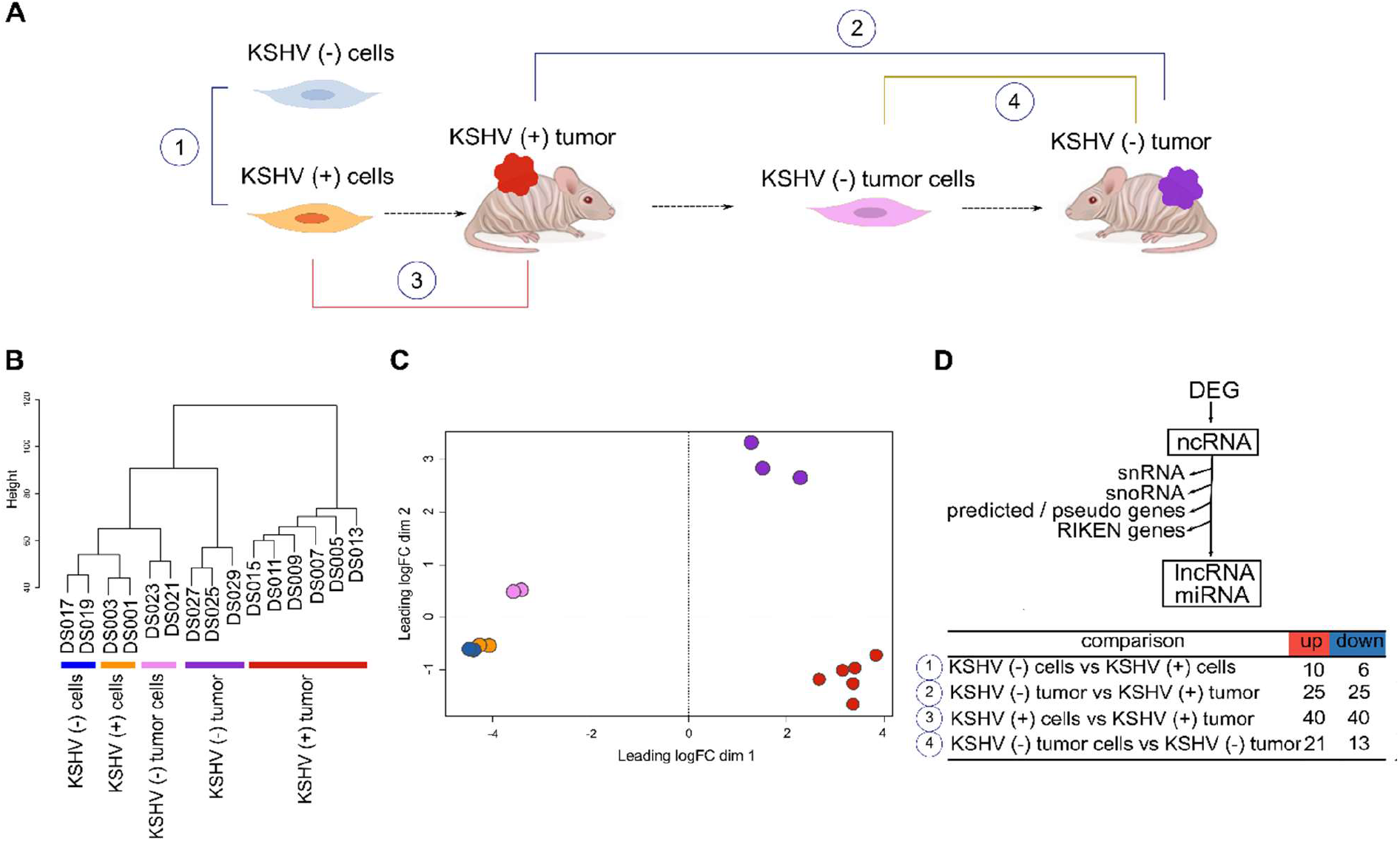
Genome-wide analysis of Non-coding RNAs in a cell and animal model of Kaposi’s Sarcoma. A) Schematic representation of the mouse-KS cell and tumor model. B) Unsupervised clustering of the host ncRNA transcriptome. C) Multidimensional scaling plot of the host ncRNAs showing the distance of each sample from each other determined by their leading log Fold Change (FC). D) Workflow analysis and number of DE lncRNAs in key biological comparisons that were detected by RNA-sequencing analysis of: two KSHV (+) cells, two KSHV (-) cells, six KSHV (+) tumors, two KSHV (-) tumor cells and three KSHV (-) tumors.

Unsupervised clustering (Figure 1B) and Multidimensional plot (Figure 1C) of the host ncRNAs shows how KSHV status and tissue type cluster with each other. As was previously reported based on mRNA proliles, in vitro and in vivo models clustered separately [Naipauer et al., 2020].

To identify changes in host lncRNAs expression profile we analyzed the number of differentially expressed (DE; FC>1.5, pvalue <0.05) lncRNAs in key biological comparisons that were detected by RNA-sequencing analysis of: two KSHV (+) cells, two KSHV (-) cells, six KSHV (+) tumors, two KSHV (-) tumor cells and three KSHV (-) tumors (S1 Table). This mouse model allows for unique experimental comparisons in the same cell and KS-like mouse tumor types: 1) KSHV (-) cells versus KSHV (+) cells can be used to study KSHV mediated effects *in vitro*, 2) KSHV (-) tumors versus KSHV (+) tumors can be used to dissect the role of ncRNAs in tumorigenesis by comparing tumors driven by KSHV versus tumors driven by host mutations, 3) KSHV (+) cells grown *in vitro* and in tumors can be used to study *in vitro* versus *in vivo* variations induced by micro-environmental cues, and 4) KSHV (-) tumor cells versus KSHV (-) tumors can be used to study *in vitro* versus *in vivo* variations in the absence of KSHV [Naipauer et al., 2020]. We first analyzed lncRNAs expression in these comparisons and found that the highest number of DE lncRNAs was observed in KSHV (+) tumors in both comparisons versus KSHV (-) tumors and versus KSHV (+) cells (Figure 1D; S1 Table).

### 2- Identification of relevant lncRNAs in KSHV (+) tumors

We performed heat map representations of all or top-50 DE lncRNAs -according to each comparison-for all the 4 biological relevant comparisons mentioned previously (Figure 2A). To select and further evaluate relevant KSHV-associated lncRNAs we searched for the common up-modulated lncRNAs in KSHV (+) tumors versus the different comparisons (Figure 2B). Of the 10 lncRNAs up-modulated in KSHV (+) cells compared to KSHV (-) cells, 3 lncRNAs (*Malat1, Neat1* and *Kcnq1ot1*) were also up modulated in the comparison of KSHV (+) tumors versus KSHV (-) tumors (Figure 2B, top panel). In addition, of the 40 up-modulated lncRNAs in KSHV (+) tumors versus KSHV (+) cells, 18 were common to the 25 up-modulated lncRNAs in the comparison between KSHV (+) tumors and KSHV (-) tumors (Figure 2B, bottom panel). These 18 genes included lncRNAs such as *Malat1, H19, Meg3, Neat1, Dio3os, Miat, Mirg*, and *Rian*, but also the miRNA genes *Mir140, Mir142, Mir27b* and *Mir378b*, among others (Figure 2B). Eventually, the analysis allowed us to select four lncRNAs with a very interesting pattern of expression through the different biological relevant comparisons (*Malat1, Neat1, H19 and MEG3*). *Malat1* and *Neat1* are upregulated in KSHV (+) cells and KSHV (+) tumors when compare with their KSHV (-) counterparts, suggesting a KSHV-dependent upregulation of these lncRNAs (Figure 2C). *H19* and *MEG3* are upregulated during the transition *in vitro* to *in vivo* in the presence of KSHV (KSHV (+) tumors versus KSHV (+) cells), but in this same transition in the absence of KSHV these lncRNAs are not upregulated (KSHV (-) tumors versus KSHV (-) tumor cells). This pattern of expression indicates a KSHV-dependent regulation of these lncRNAs during this transition induced by environmental cues (Figure 2C).

**Figure 2.**
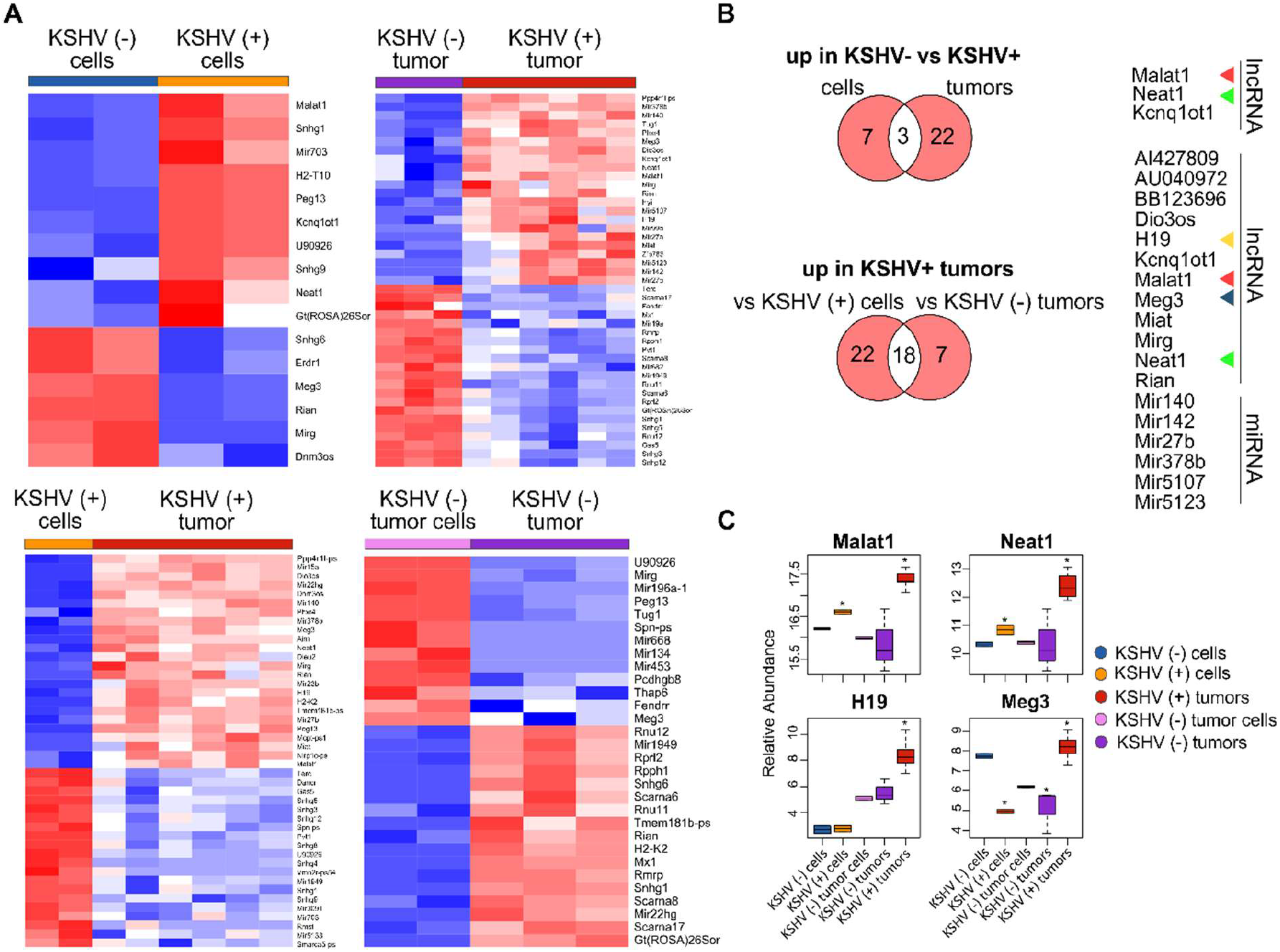
Host lncRNAs expression. A) Heat maps for fold change expression of host lncRNAs based on analysis of RNA sequencing data, all or the top 25 upregulated (red) and top 25 downregulated (blue) DE lncRNAs are shown in each comparison. B) Venn diagrams showing upregulated host lncRNAs common in KSHV (+) cells and tumors versus KSHV (-) cells and tumors (top), and upregulated host lncRNAs common in KSHV (+) tumors versus KSHV (+) cells and KSHV (+) tumors versus KSHV (-) tumors (bottom). C) Relative abundance of selected lncRNA RNAs across the different steps of the mouse-KS cell and tumor model.

### 3- Pathway analysis of the lncRNAs, reveals KSHV closely related bioprocesses

To contextualize the selected lncRNAs into functional processes, we employed LncRNA2Target v2.0 and LncTarD databases [Cheng et al., 2019; Zhao et al., 2020] as resources of lncRNA-target relationships. Since the four selected lncRNAs have been studied more extensively in humans than in mice we searched for their experimentally validated targets (EVT) (S2 Table). Functional enrichment analysis (KEGG) of the resulting lists of genes revealed several related pathways common to the four lncRNAs, mainly cancer-related pathways and bioprocesses associated with viral diseases (S1 Figure; S2 Table). Interestingly, *KSHV infection* and *MicroRNAs in cancer* were the common signature of the 4 lncRNAs (S1 Figure).

Next, we established a list of the total human target genes contributed by the 4 lncRNAs and looked for their homologues among the DE host genes previously obtained across the different comparisons of our model (Figure 3A; S3 Table). Figure 3, panels B to D, shows the chord plots illustrating the biological process terms and the target genes of the four lncRNAs contributing to that enrichment arranged in order of their expression level in the corresponding comparisons (S3 Table). Processes such as *Integrins in Angiogenesis (*with the genes *Spp1, Vegfa, Fn1, Kdr, Igf1r), Signaling by PDGF* (*Vegfa, Kdr, Cdkn1a, Igf1r*), *HIF-1 signaling (Vegfa, Hif1A, Stat3, Il6, Cdkn1a, Pik3r1, Igf1r*), *MicroRNAs in cancer* (*Dicer1, Zeb1, Zeb2, etc*.) and *KSHV infection* (*Fgf2, Hif1a, Stat3, Il6, Pik3r1, Jak2, Rb1, Jun*) were overrepresented by upregulated target genes in KSHV (+) tumors compared with KSHV (-) tumors. *Apoptosis* (*Mdm2, Bax, Myc, Casp3*) and *p53 activity regulation (Mdm2, Bax, Casp3, Pcna)* were instead associated with downregulated genes in the KSHV-bearing tumors (Figure 3B; S3 Table). Similar findings were observed in the comparison KSHV (+) cells versus KSHV (+) tumors (*in vitro* to *in vivo* transition), with the particular contribution of upregulated target genes associated with *Extracellular Matrix Organization* and *Activation of Matrix Metalloproteinases (MMP)*, represented by *Mmp2, Mmp9, Mmp13 and Mmp14* (Figure 3C; S3 Table). Also, pathways of DNA integrity control and cell cycle checkpoints, such as *Tp53 network, MicroRNA regulation of DDR* and *G1 to S cell cycle control* were revealed in this comparison, represented by the downregulated genes *E2f1, Bax, Dnmt1, Cdkn1a, Myc* or *Mdm2*. Interestingly, in the transition *in vitro* to *in vivo* but in the absence of KSHV we found processes related to the terms *KSHV infection* (*Ccnd1, Ctnnb1, Map2k2, Stat3, Pik3r1, Jun, Erbb2, Igf1r*) and *MicroRNAs in cancer* (*Dicer1, Ccnd1, Ctnnb1, Sp1*, etc.) associated with down-modulated genes in KSHV (-) tumors (Figure 3D; S3 Table), in contrast to that observed in the KSHV-dependent transition (Figure 3C).

**Figure 3:**
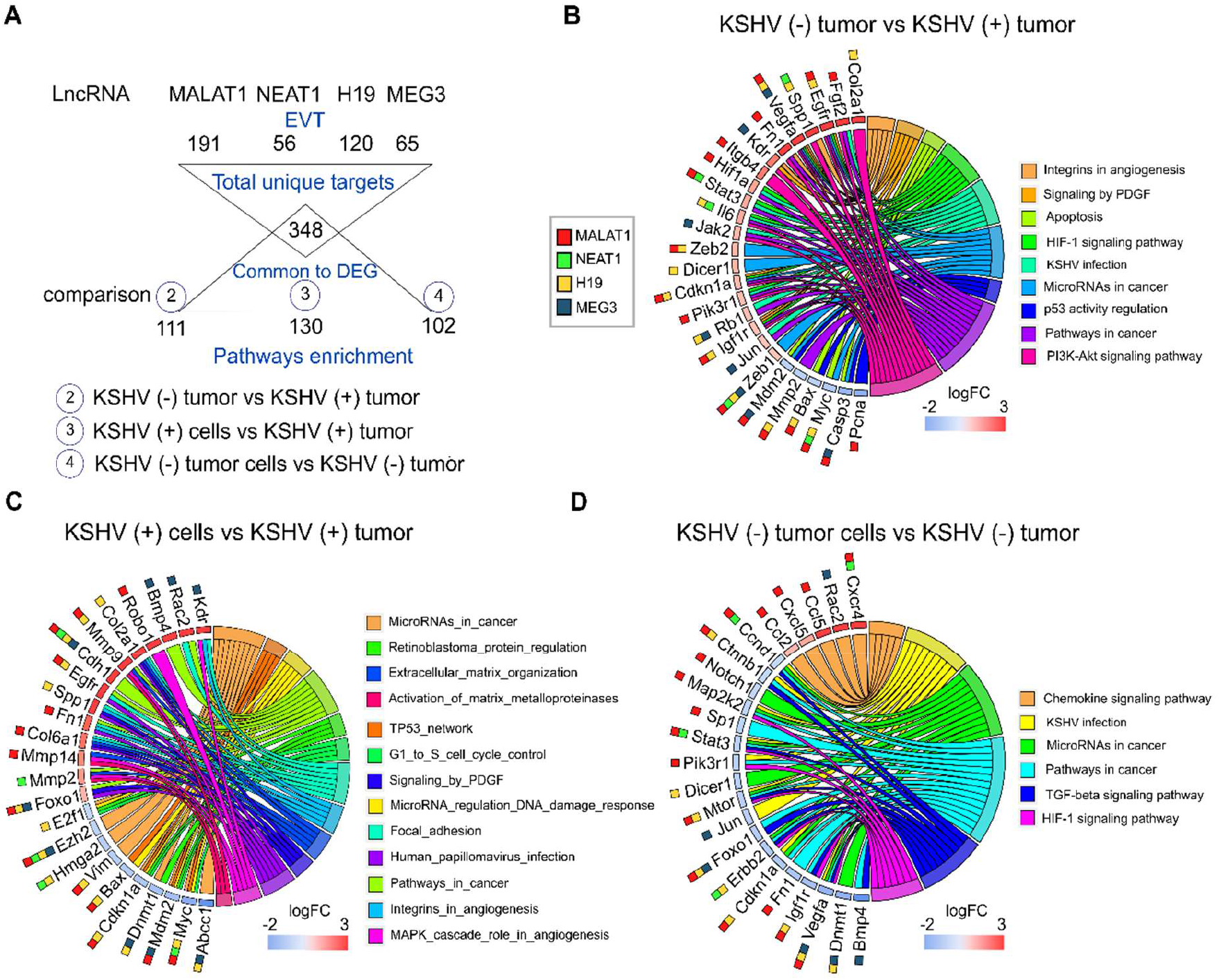
Pathway analysis of selected DE lncRNAs and their EVT genes. A) Schematic representation of the EVT genes of MALAT1, NEAT1, H19 and MEG3 that were correlated with gene expression in the RNA-sequencing analysis. B-D) Chord plot illustrating the GO biological process terms and the target genes contributing to that enrichment arranged in order of their expression level in KSHV (-) tumors versus KSHV (+) tumors (B), KSHV (+) cells versus KSHV (+) tumors (C) and KSHV (-) tumor cells versus KSHV (-) tumors (D). The corresponding lncRNAs are indicated with color boxes besides each target gene.

Taking together, the integrative in-silico analysis of the lncRNAs-EVT and their associated pathways, with the host transcriptome derived from our model, reveals that the upregulation of *Malat1, Neat1, H19* and *Meg3* in KSHV (+) tumors would contribute with a network of gene-pathways closely related with KSHV oncogenesis.

### 4- KSHV-dependent *in vitro* to *in vivo* transition is defined by a significant up-regulation of host miRNAs

LncRN6As have been demonstrated to regulate gene expression by various mechanisms, including epigenetic modifications, lncRNA-miRNA specific interactions, and lncRNAs as miRNA precursors. Our previous approach showed clear relationship among the 4 selected lncRNAs and miRNAs in cancer. Therefore, we performed small-RNA sequencing on the samples obtained from our model to identify host DE miRNAs. Next, we conducted an integrated bioinformatics workflow to elucidate relevant networks of lncRNA-miRNA-mRNA during KSHV tumorigenesis.

Unsupervised analysis of 14 samples based on miRNAs expression profiles shows how they cluster together in an unsupervised way according to their predefined features (Figure 4A). Interestingly, the samples cluster in the same pattern as when the analysis was made for lncRNAs (Figure 1B) and also for all host genes in our previous study [Naipauer et al., 2020]. The distance among groups is reflexed in the number of DE miRNAs (Figure 4B; S4 Table). Remarkably, the higher proportion of upregulated miRNAs was observed in KSHV (+) tumors (95% of DE miRNAs) compared with KSHV (-) tumors, while downregulated miRNAs were more prevalent in KSHV (-) tumors (90% of DE miRNAs) compared with KSHV (-) tumor cells (Figure 4B; S4 Table). This result is consistent with that previously described in which the term *MicroRNAs in cancer* was associated with upregulated genes in KSHV (+) tumors and down-regulated genes in KSHV (-) tumors (Figure 2B-C). Such difference could be partly explained by *Dicer1*, a master regulator of miRNA biosynthesis, which is in turn linked to the lncRNA H19 (Figure 3B and D). We performed heat map representations of all or top-50 DE microRNAs -according to each comparison-for all the 4 biological relevant comparisons mentioned previously (Figure 4C).

**Figure 4.**
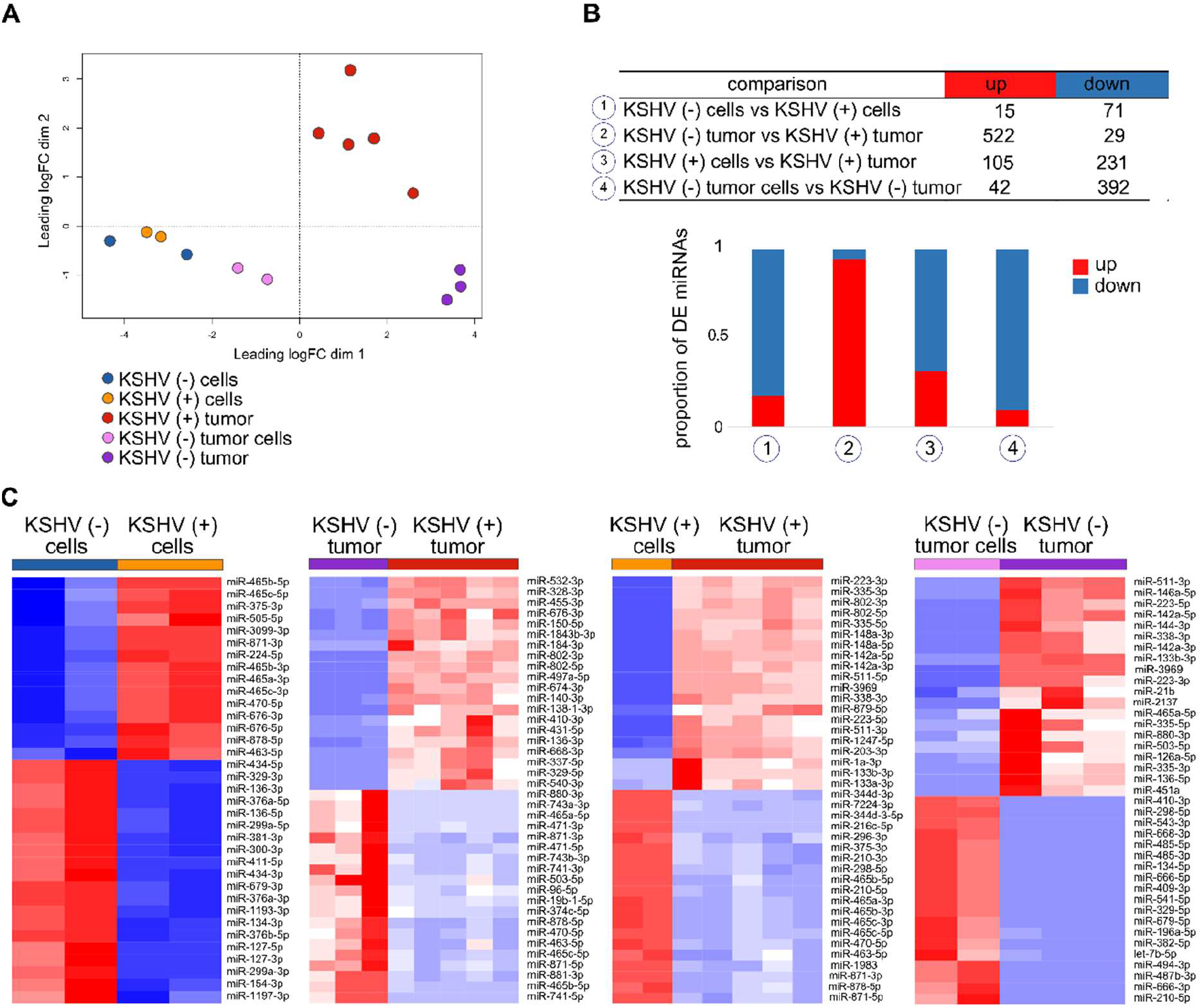
Host miRNAs expression. A) Multidimensional scaling plot of the host miRNAs showing the distance of each sample from each other determined by their leading logFC. B) Number of DE miRNAs in key biological comparisons that were detected by small RNA-sequencing analysis of: two KSHV (+) cells, two KSHV (-) cells, six KSHV (+) tumors, two KSHV (-) tumor cells and three KSHV (-) tumors. C) Heat maps for fold change expression of host miRNAs based on analysis of small RNA sequencing data, only top 20 upregulated and top 20 downregulated DE host miRNAs are shown in each comparison.

### 5- Differentially expressed miRNAs regulate gene targets related to KSHV affected biological processes

Mature miRNAs regulate gene expression at the posttranscriptional level via partial base-pairing with their target mRNAs. Such interaction leads to mRNA degradation and/or translational inhibition, causing the downregulation of proteins encoded by the miRNA-targeted mRNAs, a biological phenomenon termed RNA interference (RNAi) [Valencia-Sanchez et al., 2006]. *In silico*-based functional analysis of miRNAs usually consists of miRNA target prediction and functional enrichment analysis of miRNA targets.

To identify the experimentally supported targets from our previous published work [Naipauer et al., 2020] for the DE miRNAs identified in this study we employed DIANA TARBASE v8 [Karagkouni et al., 2018]. Next, we selected those targets whose expression antagonizes with that of its miRNA in the corresponding comparison (Figure 5A; S5 Table). As we mentioned before, most of DE miRNAs in the KSHV (-) tumors versus KSHV (+) tumors were upregulated in KSHV (+) tumors, thus their corresponding targets were downregulated in the same group. Pathways analysis of these downregulated genes indicated enrichment in: *P53 signaling pathway (Bax, Gorab, Ccng1*, Rrrm2, etc.), *Spliceosome (Tra2a, Tra2b, Srsf10, Snrpb, Snrpb2*, etc.), *E2F transcription factor (E2f6, E2f7, Rrm1, Rbl1, etc*.) and *Cell cycle (Xpo1, Nedd1, Zwint, Psma1, Psma3, etc*.), among others (Figure 5B, S5 Table).

**Figure 5.**
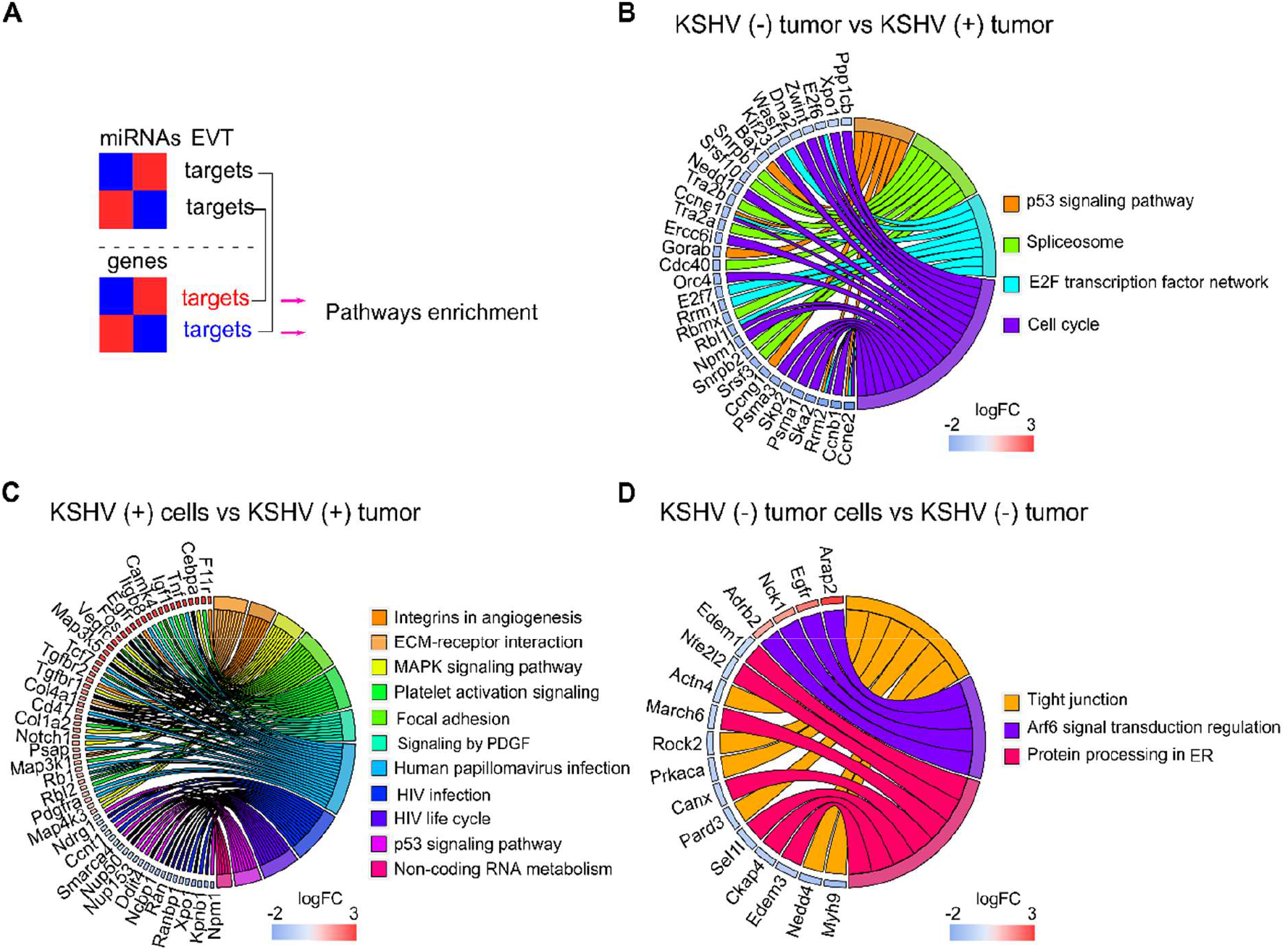
Pathway analysis of DE miRNAs and their experimentally validated target (EVT) genes. A) Schematic representation of the miRNA–mRNA pairs with significant (p < 0.05) antagonistic expression. B-D) Chord plot illustrating the GO biological process terms and the target genes contributing to that enrichment arranged in order of their expression level in KSHV (-) tumors versus KSHV (+) tumors (B), KSHV (+) cells versus KSHV (+) tumors; for a better visualization only a fraction of the genes corresponding to the plot is shown. The full list is available in S5 Table (C) and KSHV (-) tumor cells versus KSHV (-) tumors (D).

In the *in vitro* to *in vivo* transition, with a more proportional distribution of DE miRNAs, upregulated and downregulated target genes were consequently identified, which provided greater enrichment of bioprocesses closely related to the obtained with the lncRNA targets in the same comparison. As can be seen in the chord plot of Figure 5C, upregulated target genes, in the upper half of the circle, are significantly associated with processes such as *Integrins in Angiogenesis* (*Col1a12, Col4a1, Col6a2, Itgb3*, etc.), *ECM-receptor interaction* (*Itga4, Itgb3, Itgb8, Sdc1, Col1a2, etc*.*), MAPK signaling pathway* (*Rps6ka3, Cebpa, Map3k1, Map314*, etc.), *Platelet activation signaling* (*Vav3, App, Fga, Col1a2, Tgfb3*, etc.) and *Signaling by PDGF* (*Pdgfra, Col4a1, Col4a2, Col6a2, Camk4, Foxo1*, etc.). Meanwhile, in the lower half of the plot, the viral infections related processes *HPV* (*Tcf7, Fzd4, Itga4, Tcf7, Tnf*, etc.) and *HIV* (*Xpo1, Npm1, Nup50, Nup153, Nup160, Nup205*, etc.), and the *p53 signaling* (*Dusp5, Ddit4, Ccng1*, etc.) are over-represented by downregulated genes (Figure 5C, S5 Table). Within the latter, it is worth highlighting the presence of numerous genes related to the nuclear export machinery (*Ranbp1, Ran, Xpo1, Nup50, Nup153, Kpnb1, Ncbp1*). Lastly, in the *in vitro* to *in vivo* transition in the absence of KSHV, fewer terms were significantly over-represented by the miRNAs target genes. Among them highlights *Tight junctions*, linked to down-regulated genes, and *Arf6 signal transduction* over-represented by the up-regulated ones (Figure 5D, S5 Table).

Collectively, these results indicate that the presence of KSHV has a significant impact on the metabolism of host miRNAs, which contribute to the regulation of host genes linked to processes of angiogenesis, ECM, transcriptional metabolism, viral infections and cell cycle control, mainly.

### 6- KSHV miRNAs expression in mouse KSHV (+) tumors

To study the relevance of KSHV miRNA expression in KSHV tumorigenesis we used the small-RNA sequencing data of read counts to analyze the relative expression between miRNAs in KSHV (+) tumors (Figure 6A). The ten most frequent microRNAs in KSHV (+) tumors were K12-4-3p, K12-3-5p, K12-8-3p, K12-10a-3p, K12-2-5p, K12-7-3p, K12-4-5p, K12-1-5p, K12-11-3p, K12-3-3p representing 97% of the counts detected for viral microRNAs in KSHV (+) tumors (S6 Table).

**Figure 6.**
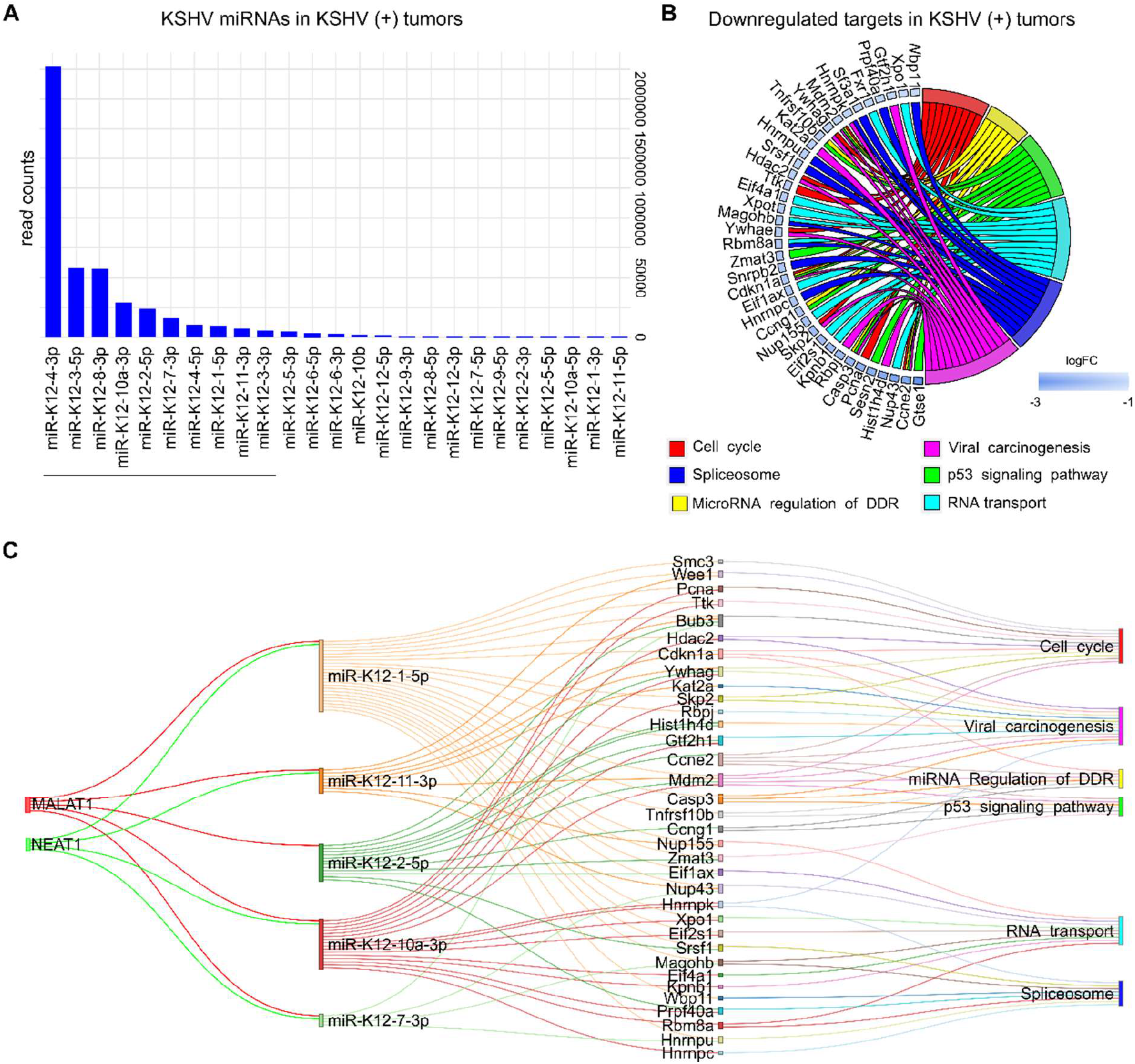
KSHV miRNAs expression analysis in KSHV (+) tumors. A) Bar plot of KSHV miRNAs relative abundance by showing counts in KSHV (+) tumors. Underlined are shown the top ten most frequent miRNAs B) Pathway analysis showing host downregulated target genes of the ten KSHV miRNAs most abundant in KSHV (+) tumors. C) Network relationship among lncRNAs, KSHV microRNAs, mRNAs and their enriched pathways.

Following the same criteria used for host ncRNAs, we searched for KSHV miRNA targets. To do this, we considered the top 10 most frequent KSHV miRNAs (Figure 6a). We used *Tarbase V8* database [Karagkouni et al., 2018], and obtained a list of 2168 human experimentally supported gene targets (S6 Table). Next, we looked for their homologues in mice, which were downregulated in KSHV (+) tumors in comparisons 2 and 3 (Figure 1A). A total of 220 genes were obtained for which functional enrichment was performed (S6 Table). Interestingly, once again, processes closely related to those previously found for host ncRNAs were obtained (Figure 6B): *Cell cycle* (*Ccne2, Cdkn1a, Hdac2, Mdm2, Pcna*, etc.); *Spliceosome* (*Hnrnpc, Hnrnpk, Hnrnpu, Magohb, Prpf40a, Sf3a1, Snrpb2, Srsf1*, etc.); *miRNA regulation of DDR* (*Casp3, Ccne2, Ccng1, Cdkn1a, Mdm2, Tnfrsf10b*); *Viral carcinogenesis* (*Casp3, Ccne2, Cdkn1, Gtf2h1, Hdac2, Hist1h4d, Kat2a, Mdm2*, etc.), *p53 signaling* (*Casp3, Ccne2, Ccng1, Cdkn1a, Gtse1, Mdm2, Sesn2, Tnfrsf10b*, etc.) or *RNA transport* (*Eif1ax, Eif2s1, Eif4a1, Fxr1, Kpnb1, Magohb, Nup155, Nup43, Xpo1, Xpot*).

### 7- Identification of a lncRNA-miRNA-mRNA interaction network involved with KSHV tumorigenesis

LncRNAs can also serve as regulatory elements of the RNAi pathway [Statello et al., 2020]. Indeed, host lncRNA transcripts are involved not only with the maturation of miRNA transcripts but also they may interfere with miRNA induced translation inhibition, thus acting as competing endogenous RNAs (ceRNAs), or “sponge RNAs” [Statello et al., 2020]. Such lncRNA-miRNA associations allow for a fine tuning of gene expression regulation. Therefore, dysregulation of the lncRNA-miRNA balance could contribute to the onset of KSHV pathogenesis.

To identify relevant pairs of lncRNA-miRNA in our model, we used DIANA-LncBase v3 [Karagkouni et al., 2020]. For each of the four lncRNAs we searched for their highly confident experimentally supported viral and host miRNA targets, derived from high-throughput methodologies, which were in turn DE in the corresponding comparison.

Within the ten most abundant KSHV miRNAs, we found that five of them have been associated with the human lncRNAs *MALAT1* and *NEAT1*. When analyzing the targets (downregulated in KSHV + tumors) of these five miRNAs, we observed that they share most of the genes obtained with the ten miRNAs, which is therefore reflected in the same enriched pathways (Figure 6C; S7 Table).

Using the same approach, we proceeded with the analyses for the lncRNA-host miRNAs associations. For the *in vitro to in vivo* transition dependent of KSHV, 31 miRNAs accomplished the criteria, of which 23 were upregulated and 8 were downregulated in KSHV (+) tumors (Figure 7; S7 Table). The highest contribution was made by *Malat1* (29 miRNAs) and *Neat1* (21 miRNAs), followed by *Meg3* (11 miRNAs) and *H19* (8 miRNAs). Among the downregulated miRNAs highlights the members of the miR17-92 family: miR-17-5p, miR-19a-3p, miR-20a-5p and miR-92a-3p. Their respective targets are represented by genes such as *Egfr, Foxo1, Pdgfra, Rb1, Igf1, Map3k1*, etc. all upregulated in KSHV (+) tumors (Figure 7). Other relevant downregulated miRNAs were miR-128-3p and miR-155-5p, which target multiple common genes. On the other hand, up-modulated miRNAs were linked mostly to *Malat1* and *Neat1*. Remarkably, among them are miR27-b-3p, miR-140-3p, miR-142-3p and miR-142-5p, whose gene precursors were also found up-modulated in KSHV (+) tumors (Figure 2B). As can be seen in Figure 7 the functional analysis that arose from the lncRNA-miRNA-mRNA triad shows that the pathways are arranged in an unsupervised way in three main clusters. The *MAPK signaling* together with *Pathways in cancer* would make up the 1st group, over-represented by the upregulated target genes contributed mainly by miR-671-5p, miR-128a-3p, miR-155-5p, and let 7e-5p, as well as the miRNAs of the miR17-92 cluster. A second group is integrated by processes related to *Viral infection* (HPV infection), *Matrix organization* and *Angiogenesis*, represented by genes contributed by the miRNAs of the miR17-92 cluster, let-7d-5p and miR-124-3p, along with others (Figure 7). A third group would be made up of the pathways *HIV life cycle* and *p53 signaling*, over-represented by negatively regulated genes, targets of the miRNAs miR-27b-3p, miR-101a-3p, miR-140-3p and miR-142, among others (Figure7).

**Figure 7.**
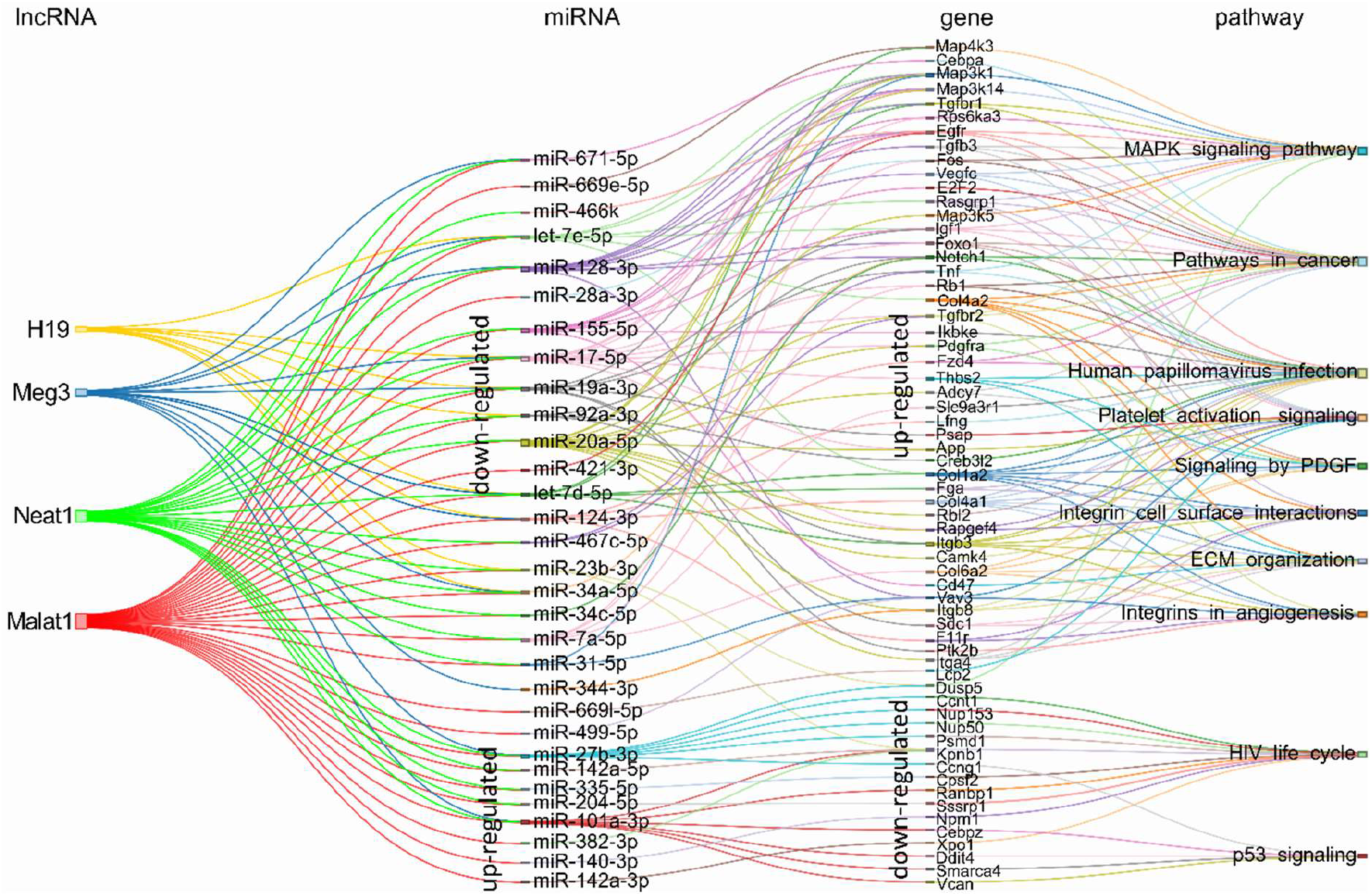
Schematic representation of the experimentally supported triad lncRNA-miRNA-mRNA and the pathways in which the later are involved (b), for the KSHV-dependent *in vitro* to *in vivo* transition. The expression status in KSHV (+) tumors is indicated for miRNAs and their respective targets.

For the comparison KSHV (-) tumors versus KSHV (+) tumors, we obtained a network of the four lncRNAs targeting 26 miRNAs all upregulated in KSHV (+) tumors with their corresponding downregulated target genes (Figure 8; S7 Table). It is evident a shift in the expression of specific miRNAs, such as let-7e-5p, let-7d-5p, miR-123-3p and miR-31-5p, compared to that observed in the KSHV dependent transition. Other relevant miRNAs that appear are miR-26b-5p, miR-181 (with its variants a, b and c), miR-378-3p and miR-381-3p. Here again, the presence of miR-140-3p and miR-378-3p correlates with their respective immature precursors that had been identified previously as upregulated along with the lncRNAs (Figure 3B). By analyzing the mRNA targets of the miRNA signature, previously identified as downregulated in KSHV (+) tumors, we obtained a relatively small group of genes that function in two major related processes: the regulation of cell cycle control (*G1 to S cycle control, p53 activity regulation, MicroRNA regulation of DDR*) and the transcription machinery, with the pre-mRNA splicing machinery (*Spliceosome)* and the *E2F transcription factor network* (Figure 8). Remarkably, this functional pattern resembles that observed with the KSHV miRNAs (Figure 6).

**Figure 8.**
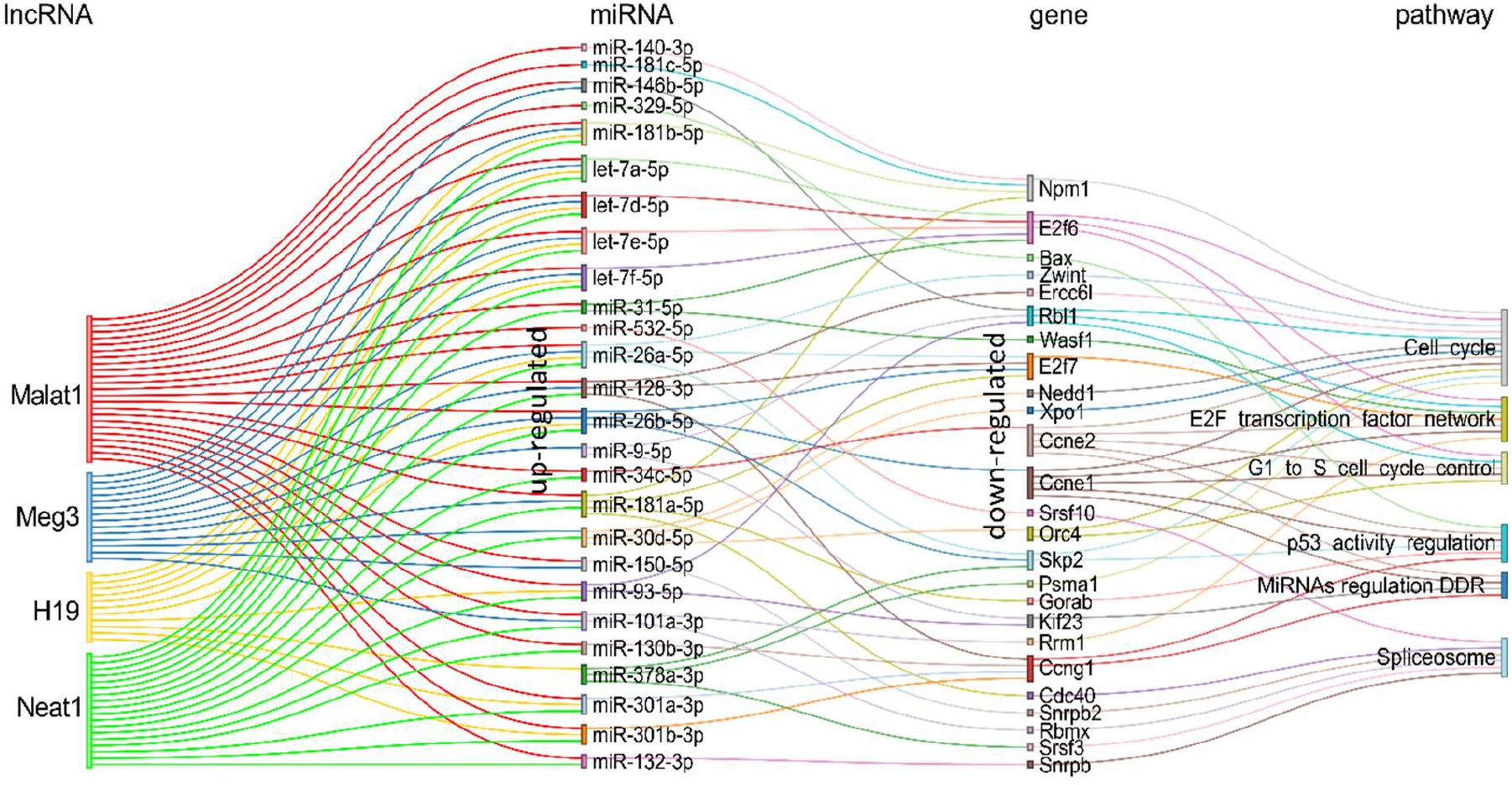
Schematic representation of the experimentally supported triad lncRNA-miRNA-gene (a) and the pathways in which the later are involved (b), for the comparison. KSHV (-) tumor vs KSHV (+) tumor. Pairs lncRNA-miRNA were identified only for the up-modulated miRNAs.

Collectively, our analysis revealed a functional network of lncRNA-miRNA-mRNA in a KSHV animal model.

### 8- Gene signatures used to identify drug-associated genes or networks

KS remains potentially life threatening for patients with advanced or ART resistant disease, where systemic therapy is indicated and three FDA-approved agents that include liposomal anthracyclines are available [Cesarman et al., 2019; Casper et al., 2011; Sullivan et al., 2008]. Despite the effectiveness of these agents, most patients progress within six to seven months of treatment and require additional therapy [Nguyen et al., 2008]. Therefore, there is a need to develop alternative strategies. Identifying drugs or clinical candidates that synergize with the current KS frontline therapeutic approaches has immediate translational potential that would be realized in a clinical trial if identified drug combinations show sustained efficacy in animal models. Our animal model allowed us to develop signatures that can be used to identify druggable gene or networks defining relevant AIDS-KS therapeutic targets.

For this end we used two approaches: 1) druggable miRNAs-gene pairs, and 2) the complete signature of the lncRNA-miRNA-mRNA network for upregulated genes.

Since miRNAs can affect the expression of druggable genes eventually affecting drug efficacy, we searched for drugs for the miRNAs-down/genes-up pairs. We employed the Pharmaco-Mir Database [Rukov et al., 2014], which identifies associations of miRNAs, genes they regulate, and the drugs dependent on these genes. S8 Table summarizes the list of drugs identified for each miRNA-gene pair. Among the drugs identified in our analysis there were some used against targets in experimental KSHV models or in clinical practice: Abacavir (mir19a-TNF), Bevacizumab (miR19a-IGF1), Celecoxib (miR17-RB1), Imatinib (miR17-PDGFRA), Oxaliplatin (miR19a-IGF1), Sirolimus (miR19a-IGF1; miR20a-MAP3K5), Sunitinib (miR-128-VEGFC; miR17-PDGFRA; miR-19a-TNF; miR-20a-PDGFRA) and Thalidomide (miR19a-TNF) (Table 1, upper half of the table).

**Table 1:**
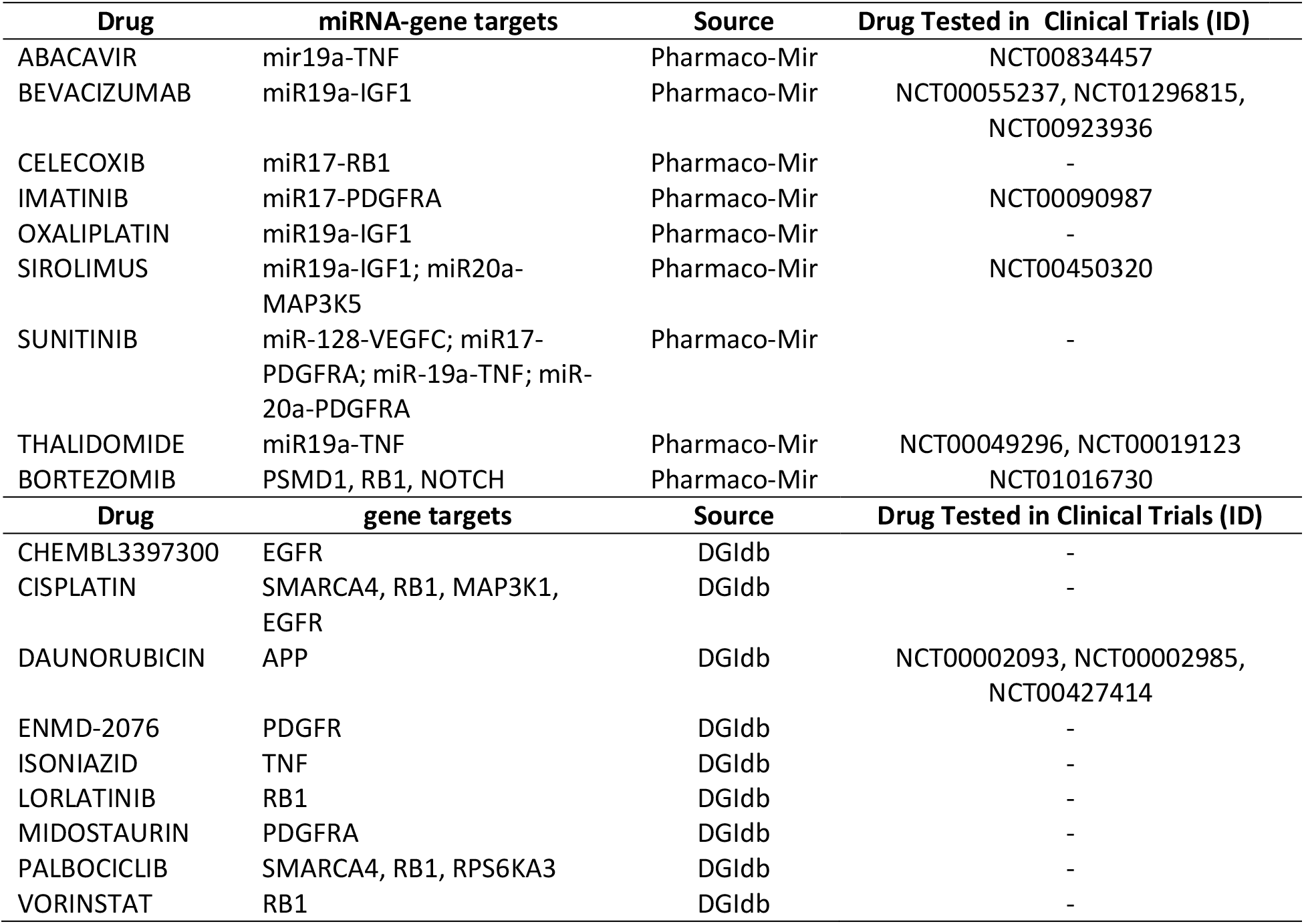
Drug-associated to miRNA-gene pairs (upper half of the table) or genes (bottom half of the table) obtained from the lncRNA-miRNA-mRNA network.

As a second approach, we used the signature of the lncRNA-miRNA-mRNA network from Figure 7 to search for drugs for the upregulated genes in the drug gene interaction database dgidb [Cotto et al., 2018]. We found, among others, chemotherapeutics agents such as Cisplatin (targeting SMARCA4, MAP3K1, RB1, EGFR, RRM1 and BAX) and Bortezomib (targeting PSMD1, RB1, NOTCH1, PSMA1 and BAX) and HDAC inhibitors, such as Vorinostat (targeting NPM1 and RB1) (Table 1, bottom half of the table). Moreover, we found kinase inhibitors such as Palbociclib (SMARCA4, RB1, RPS6KA3 and CCNE1), Midostaurin (PDGFRA) and ENMD-2076 (PDGFRA). Finally, Daunorubicin (APP) that is currently used to treat Kaposi’s Sarcoma [Petre et al., 2007]

Importantly, some of the aforementioned drugs (Abacavir, Doxorubicin, Bevacizumab, Bortezomib, Imatinib, Sirolimus and Thalidamide) have been evaluated alone or in combination with other drugs in different KS clinical trials. The description of such studies is found in S8 Table. The fact that our analyses pointed to drugs that target KS oncogenic pathways identified in the laboratory or drugs that are currently in use of being tested in AIDS-KS, reinforces the possibility of involvement of the KSHV regulated ncRNA network in viral sarcomagenesis.

## DISCUSSION

Virus-host interactions trigger a set of mechanisms that eventually affect the expression of host genes involved in the regulation of the viral replicative cycle as well as the pathogenesis of the disease [Jones et al., 2020]. Whereas dysregulation of host protein-coding genes caused by KSHV infection is well explored, host ncRNAs and KSHV dependency remains poorly characterized. Currently, miRNAs and lncRNAs are by far two of the most commonly studied ncRNA biotypes [Cech et al., 2014; Slack et al., 2019].

We have previously developed and characterized a unique multistep KSHV tumorigenesis model in which cells explanted from a KSHV (+) tumor that lose the episome can form KSHV (-) tumors driven by host mutations such as the PDGFRA-D842V [Mutlu et al., 2007; Cavallin et al., 2018]. Using NGS on this model, we interrogated the transcriptional, genetic and epigenetic (CpG island methylation) landscape upon KSHV tumor formation and upon KSHV-loss in cells and tumors [Naipauer et al., 2020]. In such study, we focused on the host and virus coding genes. Therefore, taking advantage of the model and the RNA-sequencing technology, we decided -for this study-to explore the transcriptional consequences of KSHV tumorigenesis on the ncRNAs setting, with the aim of identifying a functional interplay between lncRNAs and miRNAs dependent of KSHV.

Here we identified four relevant lncRNAs upregulated in KSHV (+) tumors: Malat1, Neat1, H19 and Meg3. Accumulating evidence has shown that lncRNA exert its functions by regulating the expression of target genes. As a first approach, using databases that collects all lncRNA–target relationships confirmed by binding experimental technologies, we searched for the target genes for the human homologues of each of the selected lncRNAs. In addition to having common target genes, pathway analysis showed that the 4 lncRNAs also share common related processes, mainly associated with cancer and viral infections. Interestingly, *KSHV infection* and *MicroRNAs in cancer* were among the common over-represented terms.

Next, we interrogated the transcriptome of our model to identify the 4-lncRNAs common targets into the DEG. The integrated analysis allowed us to define a reduced group of host lncRNAs-target genes that significantly would contribute with KSHV tumorigenesis and related processes. The integration of the in silico approach of the lncRNAs-EVT and their associated pathways, with the host transcriptome derived from our model, reveals a network of gene-pathways closely related with KSHV oncogenesis: *Integrins in angiogenesis, KSHV infection, signaling by PDGF, HIF1-signaling pathway* or *MicroRNAs in cancer* were represented by upregulated genes such as *Egfr; Vegfa, Hif1a, Dicer1, Zeb1, Zeb2, Rb1 or Il6*.

In addition, one of the distinctive pathways of the *in vitro* to *in vivo* transition dependent of KSHV, provided by the lncRNA targets, was *Extracellular Matrix Organization* and *Activation of Matrix Metalloproteinases*, overrepresented by the MMPs *Mmp2, Mmp9, Mmp13* and *Mmmp14*. MMPs are associated with KS and may contribute to the mechanism of KS tumor growth. They are usually synthesized by the tumor stromal cells, including fibroblasts, myofibroblasts, inflammatory cells and endothelial cells. These components can also integrate a tumor derived from cells *in vitro*. Although the mechanism by which Malat1, Neat1 or H19 regulate the expression of MMPs is not yet clear, different studies have shown that the silencing or overexpression of these lncRNAs positively correlate with the expression of MMPs such as MMP9 or MMP2 [Zhou et al., 2015, Liu et al., 2018; Zhu et al., 2018].

*MALAT1* is perhaps the most studied lncRNA and consequently the one with the most targets. It has been shown to regulate EGFR expression promoting carcinogenesis [Zhang et al., 2019]; it has been shown to regulate endothelial cell function and vessel growth [Michalik et al., 2014]; it has been defined as a hypoxia-induced lncRNA [Kölling et al., 2018]; it modulates *ZEB1* and *ZEB2* by sponging miRNAs [Xiao et al., 2015; Chen et al., 2017]. Remarkably, *MALAT1* expression is induced by the platelet-derived growth factor BB (PDGF-BB) [Lin et al., 2019]. In a recent study, we have shown that the KSHV-ligand mediated activation of the PDGF signaling pathway is critical for KS development [Cavallin et al., 2018]. Later, we found that two PDGFs, Pdgfa, Pdgfb and their receptor Pdgfra were both hypo-methylated and up-regulated in KSHV (+) tumors [Naipauer et al., 2020]. Overall, the evidence clearly shows that *Malat1* is a key regulator of several target genes involved in KSHV-dependent signaling pathways. It remains to be determined whether Malat1 is a driver or simply a passenger of KSHV tumorigenesis.

*NEAT1*, is closely related to *MALAT1* (aka NEAT2), and both have been shown to bind multiple genomic loci on active genes, but display distinct binding patterns, suggesting independent but complementary functions [West et al., 2014]. As *MALAT1, NEAT1* is retained in the nucleus where it forms the core structural component of the paraspeckle sub-organelles. The formation of paraspeckle increases in response to viral infection or proinflammatory stimuli [Morchikh et al., 2017]. Furthermore, Viollet et al. (2017) demonstrated that *NEAT1* is upregulated in KSHV infected cells versus non-infected cells under hypoxic conditions. Our results show that *Neat1* is upregulated in KSHV-cells versus KSHV+ cells and indeed is upregulated in KHSV (+) tumors, during the *in vitro* to *in vivo* transition. On the other hand, the lncRNA target analysis showed that *Neat1* positively associates with the upregulated targets *Il6, Stat3* and *Spp1* in the KSHV (+) tumors. In this regard, *NEAT1* has been shown to strengthen IL-6 / STAT3 signaling and promote tumor growth and proliferation through nuclear trapping of mRNAs and proteins which acts as inhibitors of the IL-6 / STAT3 signaling pathway [Wang et al., 2018]. Previously, it had been demonstrated that STAT3 is activated by KSHV infection and correlates with IL6 release in dendritic cells [Santarelli et al., 2014]. In summary, these data taking together reveal a host network in which upregulation of Neat1 would favor the activation of IL6/STAT3 signaling contributing directly or indirectly to KSHV tumorigenesis.

*MEG3* is generally considered as a tumor suppressor lncRNA. In this study we found a downregulation of *Meg3* in KSHV (-) cells versus KSHV (+) cells. However, a significant increase of the lncRNA was evidenced in the *in vitro* to *in vivo* transition. Sethuraman et al (2017) showed that KSHV employs its miRNAs to target *MEG3* promoting its downmodulation to potentially contribute to sarcomagenesis. Therefore, it is possible to speculate on a downmodulation of *Meg3* by the expressed KSHV miRNAs as an early event in the viral cycle followed by an upmodulation of *Meg3* as a response of the host cell to the already triggered tumor growth.

KSHV drives latently infected cells towards proliferation by a variety of mechanisms such as interfering with *MEG3* or the p53 pathway through miRNAs or the protein LANA, respectively (Sethuraman et al., 2017; Schulz et al., 2015). In this study, we identified that p53 network would be regulated in a KSHV dependent manner by the modulation of key genes targeted by the lncRNAs, such as *Casp3, Bax, Mdm2, Cdkn1a*, or *Pcna*. Interestingly, these genes along with *E2f1* are linked to other related processes such as *G1 to S phase regulation* and *MicroRNA DDR*. KSHV needs to face various cellular defense mechanisms designed to eradicate the viral infection. One such response can include DDR response factors, which can promote an arrest in cell growth (G1-S regulation) and trigger cell death (p53 network, Apoptosis). Our findings indicate that those processes would be repressed through the downmodulation of the mention lncRNA targets in KSHV (-) tumors versus KSHV (+) tumors, as well as in the KSHV *in vitro to in vivo* transition. Remarkably, several studies have shown viruses including KSHV have developed suppressive strategies against DDR (Weitzman and FradetTurcotte, 2018; Ohsaki E and Ueda 2020). In this sense, KSHV miRNAs are relevant for protecting cells from DDR (Liu et al., 2017). In addition, cellular *lncRNAs* are important gene regulators of DDR in a process which involve essential players of miRNA biosynthesis such as DICER1 and DROSHA [Michelini et al., 2017, Statello et al., 2020]. In fact, *Dicer1* was one of the significantly upregulated target genes linked to H19 in the comparison KSHV (-) tumors versus KSHV (+) tumors. In summary, there is a complex network between KSHV and host ncRNAs that would regulate DDR factors in order to bypass cell cycle checkpoints.

miRNA analysis revealed a high proportion of upregulated host miRNAs dependent of KSHV infection. This finding led us to interrogate the functional processes associated to the miRNAs targets. Enriched terms were linked to *p53 signaling, Spliceosome* and *Cell cycle*. When evaluating the *in vitro* to *in vivo* transition which involved both up and downregulated miRNAs, processes such as *Integrins in Angiogenesis, Platelet activation*, or *signaling by PDGF* were associated to the upregulated targets, whereas *viral infection (HPV, HIV) or p53 signaling* were linked to the downregulated targets.

Furthermore, we evaluated the relevance of viral miRNAs expression in KSHV tumorigenesis. We identified a group of ten relevant miRNAs which constituted the most frequent in mouse KSHV (+) tumors. Among them highlights K12-4-3p, K12-3-5p, K12-8-3p, previously identified as highly expressed in human KS lesions [Wu et al., 2015]. Moreover, K12-4-3p, which represented 50% of the KSHV miRNAs detected in this analysis in mouse KSHV (+) tumors, was shown to be able to restore the transforming phenotype of a mutant KSHV containing a deletion of all KSHV microRNAs [Moody et al., 2013], indicating its association with cellular transformation and tumor induction. Similarly, K12-3-5p was shown to promotes cell migration and invasion of endothelial cells [Hu et al., 2015]. The functional analysis of their targets -downregulated in mouse KSHV (+) tumors - showed enrichment in processes such as *Cell cycle, Spliceosome, RNA transport, MicroRNA Regulation of DDR* and *p53 signaling*, coinciding with what was observed with the host miRNAs, which suggests that viral miRNAs might mimic cellular miRNAs. It is possible that the same targets are also relevant to the infection of human cells by KSHV (KSHV miRNA) and to KSHV pathogenesis (host miRNA) [Forte et al., 2015, Hussein et al., 2019].

Since the similarity with the one found with respect to the lncRNA targets, we decided to carry out an integration network dependent of KSHV between the 4-lncRNAs, the DE miRNAs (from virus and host) related to those lncRNAs, their validated targets, and the related processes.

The integration showed a more concise landscape of the potential relationships of lncRNA-miRNA-mRNA in a KSHV setting, in which, once more, highlights that the upregulated genes are involved in processes such as pathways in cancer and those previously closely related to KSHV tumorigenesis, including *Angiogenesis, PDGF signaling, MAPK signaling or ECM organization*. Similarly, down-modulated genes linked preferentially to *p53 signaling, Spliceosome, miRNA regulation of DDR, or RNA transport*, among others. In the latter Proteasome subunit protein *PSMD1*, nucleoporins *NUP50, NUP153*, nucleolar protein NPM1 or exportin 1 XPO1 have been shown to modulate HIV infection or other viral cycles [Gadad et al., 2011; Behrens et al., 2017; Kane et al., 2018; Rathore et al., 2020]. In addition, in a preprint article it was postulated an extensive destruction of the nuclear and nucleolar architecture during lytic reactivation of KSHV, with redistribution or degradation of proteins such as NPM1 [Atari et al., 2020]. More interestingly NPM1 (aka NPM) is a critical regulator of KSHV latency via functional interactions with v-cyclin and LANA. Strikingly, depletion of NPM in PEL cells has led to viral reactivation, and production of new infectious virus particles [Sarek et al., 2010]. On the other hand, using a model of oncogenic virus KSHV-driven cellular transformation of primary cells, Gruffaz et al. (2019) illustrate that XPO1 is a vulnerable target of cancer cells and reveal a novel mechanism for blocking cancer cell proliferation by XPO1 inhibition.

Spliceosome has been other of the relevant terms yielded by our network analysis. In the presence of KSHV, positively regulated miRNAs linked to a group of down-modulated targets closely related to the splicing machinery (*Snrpb, Snrpb2, Rbmx, Srsf3, Srsf10*). *NEAT1* and *MALAT1* were the first lncRNAs to be identified as having a relevant role in mRNA splicing in both human and mouse cells. The mechanisms by which both lncRNAs modulate splicing is extensively reviewed in Romero-Barrios et al. (2018). Remarkably, it has been postulated that *MALAT1* modulates the phosphorylation status of a pool of Serine/arginine-rich (SR) proteins (proteins involved in splicing), resulting in the mislocalization of speckle components and changes in alternative splicing of pre-mRNAs, impacting in other SR-dependent post-transcriptional regulatory mechanisms, including RNA export, NMD and translation [Tripathi et al., 2010]. In addition, cells depleted for *MALAT1* show an increased cytoplasmic pool of poly(A)+ RNA, suggesting that *MALAT1* contribute with the retention of nuclear mRNAs. Before RNAs can interact with nuclear export machinery, they must undergo processes that regulate the number of transcripts that is exported to the cytoplasm or nuclear decay pathways. KSHV manipulation of nuclear RNA regulation is one of the strategies acquired by the virus to influence the host RNAs during viral infection [Macveigh-Fierro et al., 2020]. In fact, it was very recently demonstrated that NMD pathway targets KSHV RNAs to restrict the virus [Zhao et al., 2020]. In summary, our network reveals another intricate relationship between lncRNA-miRNA-targets that can function in modulating spliceosome pathway and RNA transport during virus-host interaction.

Other relevant miRNAs that emerged from our network were members of the cluster 17-92 and the let-7 family whose multiple targets regulate different pathways associated with cancer. Moreover, miR 140-3p or miR378b also stand out, of which, like miR143-3p, their precursors were found upregulated in KSHV (+) tumors.

It has been demonstrated that miR140 in the nucleus can interact with *NEAT1*, leading to the increased *NEAT1* expression [Gernapudi et al., 2015]. Remarkably, there is another interesting link with *NEAT1*, which is p53 signaling, a frequent pathway represented in our networks. It has been shown that silencing *Neat1* in mice prevents paraspeckle formation, which sensitizes preneoplastic cells to DDR activating cell death and impairing skin tumorigenesis [Adriaens et al., 2016]. Moreover, activation of p53 stimulates the formation of NEAT1 paraspeckles, establishing a direct functional link between p53 and paraspeckle biology [Adriaens et al, 2016]. P53 regulates NEAT1 expression to stimulate paraspeckle formation and NEAT1 paraspeckles, in turn, dampen replication-associated DNA damage and p53 activation in a negative regulatory feedback [Adriaens et al., 2016]. These data indicate that upregulation of Neat1 in KSHV (+) tumors could attenuate p53 signaling network and infected cells may benefit from this situation evading the p53 checkpoint in response to DNA damage.

As mentioned before, we have used this same mECK36 tumor model to analyze the consequences of KSHV loss by comparing the mutational and methylation landscape of KSHV (+) and KSHV (-) tumors. We found that KSHV loss led to irreversible oncogenic alterations including oncogenic mutations and irreversible epigenetic alterations that were essential in driving oncogenesis in the absence of KSHV [Naipauer et al, 2020]. In contrast to these irreversible effects of KSHV tumorigenesis, the ncRNA network we describe in the present study display a high degree of plasticity and reversibility upon KSHV loss further supporting the idea that these oncogenic networks are driving tumorigenesis and are more strictly dependent on the presence of KSHV.

Finally, this ncRNA-mRNA analysis in the animal model presented here allowed us to develop signatures that can be used to identify druggable gene or networks defining relevant AIDS-KS therapeutic targets. Interestingly, among the drugs we identified are those usually used against targets in experimental KSHV models or in clinical trials: Abacavir, Bevacizumab, Bortezomib, Celecoxib, Doxorubicin, Imatinib, Oxaliplatin, Sirolimus, Sunitinib, Thalidomide and Vorinostat. Interestingly, we have previously shown a combinatory effect between Bortezomib and Vorinostat for the treatment for primary effusion lymphoma [Bhatt et al., 2013]. The fact that our analyses pointed to drugs that target KS oncogenic pathways identified in the laboratory or drugs that are currently in use of being tested in AIDS-KS further validate of bioinformatic analysis for efficient identification of druggable gene or networks defining relevant AIDS-KS therapeutic targets. It also reinforces the idea of the involvement of the KSHV regulated ncRNA network in viral sarcomagenesis.

In summary, in the present study the integration of the transcriptional analysis of ncRNAs in a KSHV model in cells and mouse tumors, with an exhaustive computational analysis of their experimentally supported targets, has allowed us to dissect a complex network that defines the main pathways involved in KSHV pathogenesis and host response. Understanding the relationships between these different RNA species will allow a better understanding of the biology of KSHV and can aid in the identification of relevant AIDS-KS druggable targets.

## METHODS

### RNA-Sequencing analysis

RNA-sequencing raw data used in the present study was obtained as previously described [Naipauer et al, 2020]. Data are available at https://www.ncbi.nlm.nih.gov/geo/, GSE144101. Briefly, paired-end sequencing using an Illumina NextSeq500 platform was used. All samples were processed in the same sequencing run of Illumina NextSeq 500 system and analyzed together with the aim to avoid the batches effect. The short sequenced reads were mapped to the mouse reference genome (GRCm38.82) by the splice junction aligner TopHat V2.1.0. Several R/Bioconductor packages to accurately calculate the gene expression abundance at the whole-genome level using the aligned records (BAM files) were used. The number of reads mapped to each gene based on the Mus musculus genome assembly GRCm38 (mm10) were counted, reported and annotated using the featureCounts package. To identify DE genes between cell lines and tumor samples, we utilized the DESeq2 package based on the normalized number of counts mapped to each gene. For ncRNA annotation we employed biomaRt package. We considered the Ensemble transcript ID, the Ensembl gene ID, the Entrezgene ID, the HGNC symbol, the Refseq ncRNA ID and the ReqSeq ncRNA predicted ID. After Deseq2 analysis on all ncRNAs, we filtered out those belonged to the following classes: small nuclear RNA (snRNA), small nucleolar RNA (snRNA), predicted and or pseudogenes, and RIKEN genes; and kept the classes lncRNA and miRNA.

### Small RNA sequencing and miRNAs analysis

Cells and tumors employed in the present study were the same as previously described (Naipauer et al., 2020). Briefly, mECK36, KSHV (+) cells were originated from frozen batches of mECK36 cells previously generated [Mutlu et al., 2007]. KSHV (+) tumors were obtained as previously shown, 1×106 KSHV (+) cells were injected subcutaneously into the flanks of nude mice and KSHV (+) tumors formed 5 weeks after injection. KSHV (-) cells were used from frozen populations of KSHV null mECK36 previously obtained [Mutlu et al., 2007]. KSHV (-) tumor cells were obtained from frozen stocks previously generated by explanted mECK36 tumor cells that have lost the Bac36-KSHV episome [Mutlu 2007]. These KSHV-negative cells were obtained from frozen stocks previously generated [Ma et al., 2013]. KSHV (-) tumors were obtained as previously shown [23], 1×106 KSHV (-) tumor cells were injected subcutaneously into the flanks of nude mice and KSHV (-) formed tumors 3 weeks after injection.

RNA was isolated and purified using RNeasy Plus Mini Kit (Qiagen, #74134) following the RNeasy MinElute Cleanup Kit (Qiagen, #74204) to separate purification of small RNA (containing miRNA) and larger RNA, the small RNA eluate is enriched in various RNAs of <200 nucleotides. A total of 15 small RNA ranged from cell lines to primary mouse tumors in the presence or absence of KSHV, were processed and sequenced on a HiSeq 2500 System (Illumina, USA). Each sample yielded on average 17 million reads, with the exception of one sample (DS016) that was excluded from the analysis for presenting a low number of total reads. Nearly all bases showed scores > Q30 for all reads. Trimmomatic was used to remove adapters and quality control was checked with FastQC. Reads were mapped to a combined mouse and KSHV genome using the bowtie aligner (ver. 1.1.1). To identify novel and known miRNAs we used miRDeep2 package (ver. 2.0.0.7). A hybrid genome of the mouse and the KSHV virus was used for all analyzes in order not to bias the mapping results for or against any of the two separate genomes. The source for all known miRNAs was miRBase (ver. 21). KSHV transcriptome was analyzed using previous resources and KSHV 2.0 reference genome [Arias 2014], while Deseq2 test was employed for differential gene expression analysis of KSHV transcripts.To identify DE miRNAs across the different comparisons, we utilized the DESeq2 test based on the normalized number of counts mapped to each miRNA. For data integration and visualization of DE transcripts we used R/Bioconductor.

### Integrative computational and bioinformatics analysis

To identify experimentally validated target genes regulated by the selected lncRNAs we employed LncRNA2Targetv2.0 (http://123.59.132.21/lncrna2target) and LncTarD (http://biocc.hrbmu.edu.cn/LncTarD/) databases [Cheng 2019; Zhao 2020]. To obtain the experimentally supported targets of the DE host miRNAs identified in this study, we employed DIANA TARBASE v8 (https://carolina.imis.athena-innovation.gr/diana_tools/). For KSHV miRNAs targets we also used DIANA TARBASE v8 resource [Karagkouni 2018] To identify relevant pairs of lncRNA-miRNA in our model, we used DIANA-LncBase v3 [Karagkouni 2020]. For each of the four lncRNAs we searched for their highly confident experimentally supported viral and host miRNA targets. To identify drug-associated genes or networks we used the drug gene interaction database (DGIdb; https://www.dgidb.org/) and the miRNA Pharmacogenomics Database (Pharmaco miR; http://www.pharmaco-mir.org/) [Cotto 2018; Rukov 2014] ClinicalTrials.gov database (https://clinicaltrials.gov/) was consulted to search for all recruiting and non-recruiting studies of KS patients.

Functional enrichment analyses were performed using the ClueGo Cytoscape’s plug-in (http://www.cytoscape.org/) and the Enrichr resource (https://maayanlab.cloud/Enrichr/) based on the lists of EVT that were in turn deregulated transcripts across the different comparisons of our model. For pathways terms and annotation, we used those provided by KEGG and BioPlanet (http://tripod.nih.gov/bioplanet/; https://www.genome.jp/kegg/pathway.html).

All statistical analysis and data visualization plots were done with R/Bioconductor packages.

## Acknowledgments

We would like to thank the Oncogenomics Core Facility at the Sylvester Comprehensive Cancer Center from the University of Miami and the Laboratory Core of the Miami CFAR for performing high-throughput sequencing.

---

**S1 Figure:**
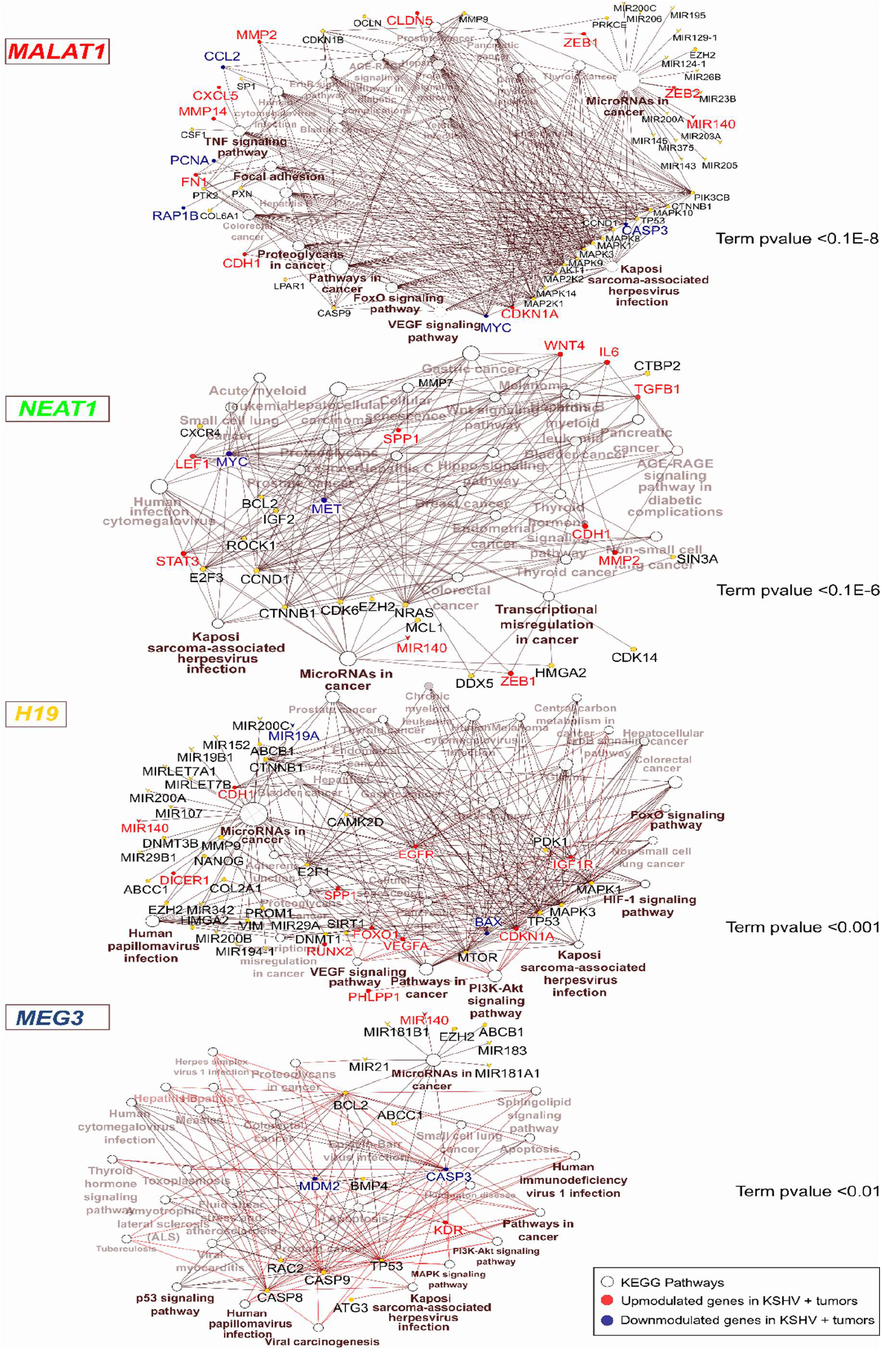
LncRNA EVT network and their related pathways. Highlighted in red and blue are the upregulated and downregulated genes (FC >1.5; pvalue <0.05) in the KSHV+ tumors of the mouse model, respectively.

**S1 Table:** DE lncRNAs in key biological comparisons detected by RNA-sequencing. Results were obtained after DeSeq2 analysis of: two KSHV (+) cells, two KSHV (-) cells, six KSHV (+) tumors, two KSHV (-) tumor cells and three KSHV (-) tumors.

**S2 Table:** Pathway analysis of the lncRNAs EVT.

**S3 Table:** Pathway analysis of the selected lncRNAs and their EVT genes DE in the corresponding comparisons.

**S4 Table:** DE miRNAs in key biological comparisons detected by small RNA-sequencing.

**S5 Table:** Pathway analysis of DE miRNAs and their EVT genes DE in the corresponding comparisons.

**S6 Table:** KSHV miRNAs analysis in KSHV (+) tumors and pathway analysis of their EVT.

**S7 Table:** lncRNA-miRNA-mRNA-Pathway networks

**S8 Table:** Drugs associated with miRNA-gene pairs obtained from network analysis.

